# Recurrent gene duplication leads to diverse repertoires of centromeric histones in Drosophila species

**DOI:** 10.1101/086942

**Authors:** Lisa E. Kursel, Harmit S. Malik

## Abstract

Despite their essential role in the process of chromosome segregation in most eukaryotes, centromeric histones show remarkable evolutionary lability. Not only have they been lost in multiple insect lineages, but they have also undergone gene duplication in multiple plant lineages. Based on detailed study of a handful of model organisms including *Drosophila* melanogaster, centromeric histone duplication is considered to be rare in animals. Using a detailed phylogenomic study, we find that *Cid*, the centromeric histone gene, has undergone four independent gene duplications during Drosophila evolution. We find duplicate *Cid* genes in *D. eugracilis* (*Cid2*), in the *montium* species subgroup (*Cid3, Cid4*) and in the entire *Drosophila* subgenus (*Cid5*). We show that Cid3, Cid4, Cid5 all localize to centromeres in their respective species. Some *Cid* duplicates are primarily expressed in the male germline. With rare exceptions, *Cid* duplicates have been strictly retained after birth, suggesting that they perform non-redundant centromeric functions, independent from the ancestral *Cid*. Indeed, each duplicate encodes a distinct N-terminal tail, which may provide the basis for distinct protein-protein interactions. Finally, we show some Cid duplicates evolve under positive selection whereas others do not. Taken together, our results support the hypothesis that *Drosophila* Cid duplicates have subfunctionalized. Thus, these gene duplications provide an unprecedented opportunity to dissect the multiple roles of centromeric histones.

**Author Summary:** Centromeres ensure faithful segregation of DNA throughout eukaryotic life, thus providing the foundation for genetic inheritance. Paradoxically, centromeric proteins evolve rapidly despite being essential in many organisms. We have previously proposed that this rapid evolution is due to genetic conflict in female meiosis in which centromere alleles of varying strength compete for inclusion in the ovum. According to this ‘centromere drive model’, essential centromeric proteins (like the centromeric histone, CenH3) must evolve rapidly to counteract driving centromeres, which are associated with reduced male fertility. A simpler way to allow for the rapid evolution of centromeric proteins without compromising their essential function would be via gene duplication. Duplication and specialization of centromeric proteins would allow one paralog to function as a drive suppressor in the male germline, while allowing the other to carry out its canonical centromeric role. Here, we present the finding of multiple *CenH3* (*Cid*) duplications in *Drosophila.* We identified four instances of Cid duplication followed by duplicate gene retention in *Drosophila.* These *Cid* duplicates were born between 20 and 40 million years ago. This finding more than doubles the number of known *CenH3* duplications in animal species and suggests that most *Drosophila* species encode two or more *Cid* paralogs, in contrast to current view that most animal species only encode a single *CenH3* gene. We show that duplicate Cid genes encode proteins that have retained the ability to localize to centromeres. We present three lines of evidence, which suggest that the multiple Cid duplications have been retained due to subfunctionalization. Based on these findings, we propose the novel hypothesis that the multiple functions carried out by CenH3 proteins*, i.e.,* meiosis, mitosis and gametic inheritance, may be inherently incompatible with one another when encoded in a single locus.

## Introduction

Centromeres are the chromosomal regions that link DNA to the spindle during cell division, thus ensuring faithful segregation of genetic material. Proper centromere function is critical for eukaryotic life. Centromeric defects can result in aneuploidy and cycles of chromosome breakage [1, 2] with catastrophic consequences for genome stability and fertility. Despite the fact that centromeres are essential for life, centromere architecture is remarkably diverse [3]. Centromeric DNA sequences [4-6] and centromeric proteins [7-9] also evolve rapidly in diverse organisms. This diversity and rapid evolution make it nearly impossible to name a single defining feature of all centromeres. However, the hallmark of many centromeres is the presence of a specialized centromeric H3 variant called CenH3 (CENP-A in mammals [10, 11], Cid in *Drosophila* [12]). Despite being essential for chromosome segregation in most eukaryotes [13-15], *CenH3* evolves rapidly [7, 16] Thus, paradoxically, proteins and DNA that mediate chromosome segregation in eukaryotes are less conserved than one would expect given their participation in an essential process. This rapid evolution despite the expectation of constraint is referred to as the ‘centromere paradox’ [17].

Genetic conflicts provide one potential explanation for the rapid evolution of centromeric DNA and proteins. In both animals and plants, the asymmetry of female meiosis provides an opportunity for centromere alleles to act selfishly to favor their own inclusion in the oocyte and subsequent passage into offspring rather than the polar body. In female meiosis, centromeric expansions [18] and differential recruitment of centromeric proteins resulting in centromere strength variation between homologs [19] may provide the molecular basis of segregation distortion. In males, however, expanded centromeres and centromere strength variation are thought to result in reduced fertility [18, 20]. This lower fertility is predicted to drive the evolution of genetic suppressors of ‘centromere drive’, including alleles of centromeric proteins with altered DNA-binding affinity. Under this model, centromeric proteins evolve rapidly in order to mitigate fitness costs associated with ‘centromere drive’ [21].

‘Centromere drive’ and its suppression provide an explanation for the rapid evolution of both centromeric DNA and centromeric proteins. However, it invokes the relentless, rapid evolution of essential proteins such as CenH3, whose mutation could be highly deleterious [13-15, 22]. A simpler way to allow for the rapid evolution of centromeric proteins without compromising their essential function would be via gene duplication. Duplication and specialization of centromeric proteins would allow one paralog to function as a drive suppressor in the male germline, while allowing the other to carry out its canonical centromeric role. Gene duplication as a way of separating functions with divergent fitness optimums has been previously invoked to explain the high frequency of duplicate gene retention, including retention of testis-expressed gene duplicates that carry out mitochondrial functions [23]. Even though both somatic and testis mitochondrial functions are similar, they have different fitness maxima, which may not be simultaneously achievable using the same set of genes. For example, the most important selective constraint shaping mitochondrial function in sperm may be the increased production of faster-swimming sperm even at the expense of a higher mutation rate. A high mitochondrial mutation rate in sperm is mitigated by the fact that sperm mitochondria are not transmitted to offspring, however such a high mutation rate would be deleterious for somatic tissues. Gene duplications allow organisms to achieve optimal mitochondrial function simultaneously in somatic tissues and testes. By the same reasoning, if a single-copy gene is incapable of achieving the multiple fitness optima that are required for multiple centromeric functions *(e.g*., mitosis versus meiosis), gene duplication could allow each duplicate to achieve optimality for different functions, thereby resolving intralocus conflict [23]. The potential for functional interrogation of intralocus conflict within *CenH3* makes the identification and study of *CenH3* duplications intriguing.

At least five independent gene duplications of *CenH3* have been described in plants [24-30]. In most cases, both protein variants are widely expressed and co-localize at centromeres during cell divisions [26, 27]. However, in barley, one *CenH3* paralog is widely expressed while the other is only expressed in embryonic and reproductive tissues [29]. In cases that have been examined closely, *CenH3* duplicates are subject to divergent selective pressures *(i.e.* one paralog evolves under positive selection but the other does not) [25, 27]. Indeed, *CenH3* duplications in *Mimulus guttatus* have been hypothesized to result from centromere-drive suppression [25].

Despite this evidence of frequent *CenH3* duplication in plants, *CenH3* duplications have been sparsely characterized in animals. So far, only three examples of *CenH3* duplications have been described: in the holocentric nematodes *Caenorhabditis elegans* and *C. remanei [31, 32]* (thought to be independent duplications) and in Bovidae (including cows) [33]. Detailed studies have only been performed on the *CenH3* duplicate in *C. elegans,* and these have yet to elucidate a clear function [32]. Similarly, only two of the ten cow *CenH3* duplicates have retained open reading frames and all cow duplicates remain poorly characterized [33]. This dearth of data has led to the suggestion that, owing to their propensity to undergo polyploidization events [34, 35], plants are more likely to retain *CenH3* duplications than animals [36].

An alternative explanation is that our perception is skewed because many systems in which *CenH3* has been extensively studied (predominant mammalian systems, such as mice and humans, and model organisms like *D. melanogaster)* have only one copy of *CenH3.* To address this possibility, we took advantage of the recent sequencing of high-quality genomes from multiple *Drosophila* species. These genomes are at a close enough evolutionary distance to allow inferences of gains, losses and selective constraints. Despite there being only one copy of *CenH3* in *D. melanogaster*, we were surprised to find that some *Drosophila* species had two or more copies of *CenH3.* This motivated our broader analysis of *CenH3* duplication and evolution throughout *Drosophila.* In total, we find at least four independent *Cid* duplications over *Drosophila* evolution. Cytological analyses confirm that these *Cid* duplicates encode *bona fide* centromeric proteins, two of which are expressed primarily in the male germline. Based on their retention without loss over long periods of *Drosophila* evolution, and analysis of their selective constraints, we infer that these duplicates now perform non-redundant centromeric roles, possibly as a result of subfunctionalization. Overall, this suggests that *Drosophila* species encoding a single *CenH3* gene may be in the minority. The sheer number of available *Drosophila* species and their experimental tractability make *Drosophila* an ideal system to study the evolution and functional specialization of duplicate *Cid* genes. Our results suggest the intriguing possibility that *CenH3* duplications may allow *Drosophila* species to better achieve functional optimality of multiple centromeric functions *(e.g*., mitotic cell division in somatic cells and centromere drive suppression in the male germline) than species encoding a single *CenH3* gene.

## Results

### Four *Cid duplications* in the *Drosophila* genus: ancient retention and recent recombination

Although their N-terminal tails are highly divergent, CenH3 histone fold domains (HFD, ~100 aa) are highly conserved and recognizably related to canonical H3 [37, 38]. Thus, sequence similarity searches based on either CenH3 or even canonical H3 HFDs are sufficient to identify putative *CenH3* homologs in fully sequenced genomes; inability to find homologous genes can be indicative of true absence [39]. To identify all *CenH3* homologs in *Drosophila,* we performed a tBLASTn search using both the canonical H3 and the *D. melanogaster* CenH3 (Cid) HFD as a query against 22 sequenced *Drosophila* genomes, as well as genomes from two additional dipteran species. We recorded each *Cid* gene “hit” as well as its syntenic locus in each species (Fig 1A, Table S1). Consistent with previous studies, we found no additional *Cid* genes in the *D. melanogaster* genome or in closely related species of the *melanogaster* species subgroup. In addition, we found that orthologs of the *Cid* gene in *D. melanogaster* have been preserved in their shared syntenic location in each of the *Drosophila* species we examined, except in *D. eugracilis* where it has clearly pseudogenized (Fig S1). We also found *Cid* orthologs in the shared syntenic context in a basal *Drosophila* species, *D. busckii,* as well as *Phortica variegate*, which belongs to an outgroup sister clade of *Drosophila*. Based on these findings, we conclude that an ortholog of *D. melanogaster Cid1* was present in the common ancestor of *Drosophila* in the shared syntenic location. We denote this orthologous set of genes in this shared syntenic location as *Cid1*.

**Fig 1.**
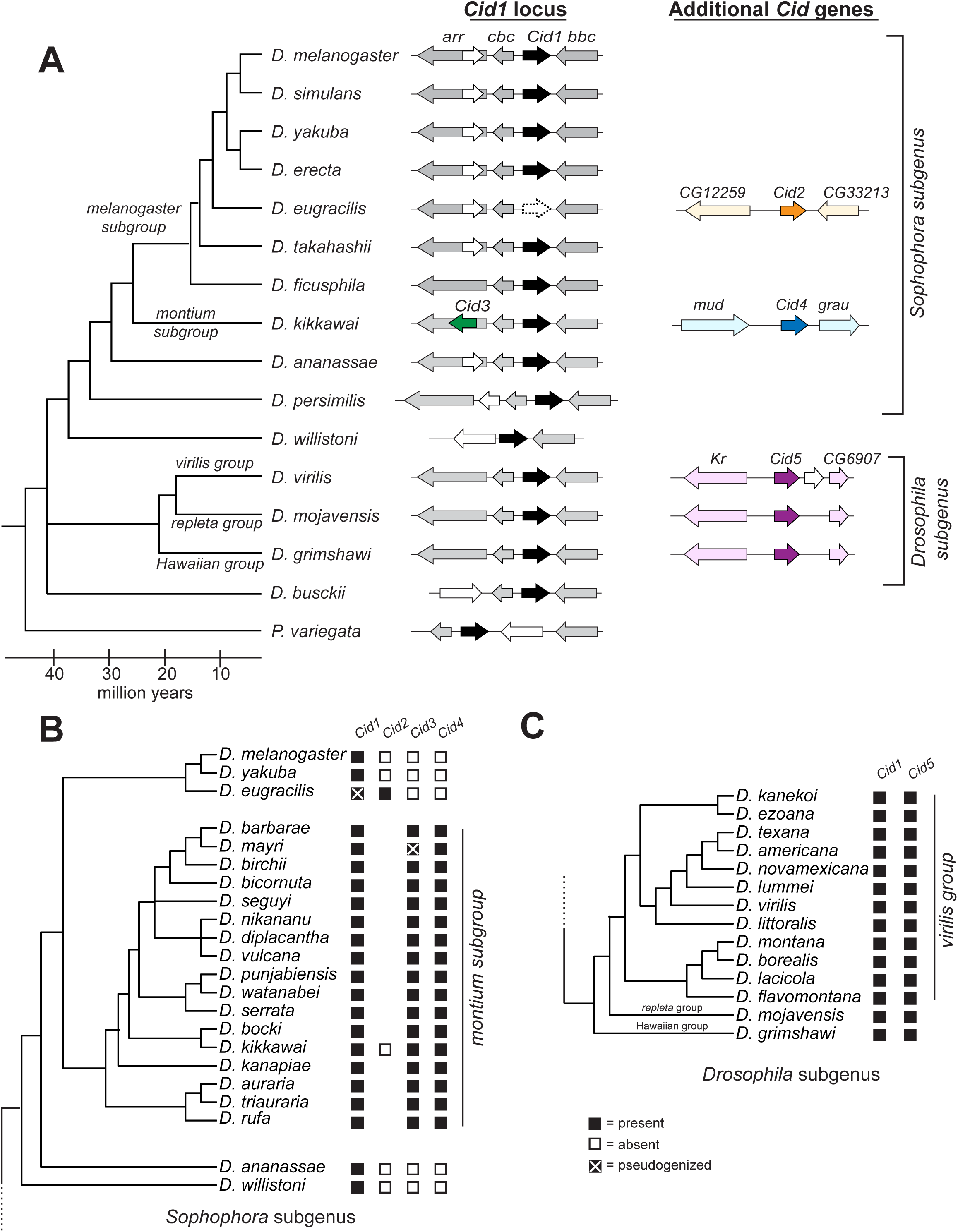
Identification of *Cid* duplication events across *Drosophila* evolution. (A) A *Drosophila* species cladogram is presented with *Phortica variegata* as an outgroup. The genomic context of representative *Cid* paralogs identified by tBLASTn using previously published genome sequences is schematized to the right of each species. *Cid1* is the ancestral locus based on its presence in almost all species, including the outgroup species *P. variegata* (black arrow, see column labeled *‘Cid1* locus’). In total, we found four *Cid* duplication events resulting in the birth of the genes *Cid2, Cid3, Cid4* and *Cid5* (see *‘Cid1* locus’ and ‘Additional *Cid* genes’ columns, dark orange, dark green, dark blue and dark purple arrows). We also found one *Cid1* pseudogene *(Cid1* locus’ column, empty arrow, dashed outline) in *D. eugracilis.* Arrows colored in a lighter version of the corresponding *Cid* gene color represent genes that define the shared syntenic locus of each paralog. White arrows represent genes that are present in a locus, but do not define the locus since they are present in fewer than 50% of the represented species. We do not provide gene names for these ‘white arrow’ genes. Genes that define each syntenic locus are named based on the *D. melanogaster* gene name. (B) Summary of *Cid* paralog presence across the *Sophophora* subgenus with an expanded *montium* subgroup. The presence (black box) or absence (white box) of each *Cid* paralog as determined by PCR and Sanger sequencing is displayed next to each species. The lack of a box means that we did not attempt to amplify the locus. *Cid1*, *Cid3* and *Cid4* were preserved in almost all *montium* subgroup species with the exception of a *Cid3* pseudogene in *Drosophila mayri* (black box with a white X). This analysis indicated that *Cid3* and *Cid4* were born 20 – 30 million years ago. (C) Summary of *Cid* paralog presence across the *Drosophila* subgenus with an expanded *virilis* group. *Cid1* and *Cid5* were completely preserved in all *virilis* group species. We conclude that *Cid5* was born 40 – 50 million years ago in the common ancestor of the *Drosophila* subgenus.

Our analysis also identified four previously undescribed *Cid* duplications in *Drosophila* (Fig 1A). The first of these was in *D. eugracilis,* which has a pseudogene at the ancestral *Cid1* shared syntenic location but also encodes a full-length *Cid* gene in a new syntenic location (Fig 1A, S1). We refer to this gene as *Cid2.* We sequenced an additional 8 strains of *D. eugracilis* to see if there were any cases of dual retention of both *Cid1* and *Cid2* in this species (Data S1). In all cases, we found that *Cid1* orthologs were pseudogenized; they all contained a two base pair deletion leading to a frame shift after the first nine amino acids and a stop codon after 12 amino acids. *D. eugracilis* represents a unique case wherein the ancestral *Cid1* was lost and replaced by a recent duplicate, *Cid2.* Based on additional sequencing (below) it remains the only case of *Cid1* loss described in *Drosophila*.

In addition to the *Cid* duplicate in *D. eugracilis,* we found two new *Cid* paralogs in *D. kikkawai*, which belongs to the *montium* subgroup of *Drosophila*. Thus, *D. kikkawai* encodes three *CenH3* genes: the ancestral *Cid1,* as well as *Cid3* and *Cid4* (Fig 1A). *Cid3* is located in close proximity to the original *Cid1* gene, whereas *Cid4* is present at a distinct syntenic location. *Cid1, Cid3* and *Cid4* are quite different from one another at the sequence level. Their N-terminal tails only share ~25% amino acid identity, whereas pairwise amino acid identity of their HFD ranges from 80% (Cid1 and Cid3) to 55% (Cid3 and Cid4) to 45% (Cid1 and Cid4). To study the age and evolutionary retention of these *Cid* paralogs, we sequenced these three syntenic loci from 16 additional species of the *montium* subgroup, for which no genomic sequences are publically available. We found that *Cid1, Cid3* and *Cid4* have been almost completely preserved in the *montium* subgroup (Fig 1B) with one exception: the *Cid3* ortholog is pseudogenized in *D. mayri* (Fig 1B, S2). Due to the lack of a complete genome sequence, we cannot rule out the possibility that *D. mayri* encodes a *Cid3*-like gene elsewhere in its genome. Based on these findings, we conclude that *Cid3* and *Cid4* were born from duplication events in the common ancestor of the *montium* subgroup at least 15 million years ago [40].

The fourth Cid duplication was found in the three species of the *Drosophila* subgenus: *D. virilis, D. mojavensis* and *D. grimshawi* (Fig 1A, ‘Additional *Cid* genes’ column). Each of these species encodes *Cid1* and *Cid5,* which have an average pairwise amino acid identity of 60% in the HFD but only 15% in the N-terminal tail. To investigate the age and evolutionary retention of *Cid1* and *Cid5,* we sequenced both genes from an additional 11 species from the *virilis* species group. We found that both *Cid1* and *Cid5* have been completely preserved (Fig 1C). Thus, we conclude that *Cid5* was born in the common ancestor of *Drosophila* subgenus at least 40 million years ago [40].

To more rigorously test the paralogy and age of the *Cid* duplicates, we performed phylogenetic analyses (Fig 2). The N-terminal tails of all the Cid proteins were too divergent to be aligned, so we built a codon-based DNA alignment of the HFD of all *Drosophila Cid* genes, including *Cid1* orthologs sequenced in a previous survey [41] (for untrimmed sequences see Data S2, for alignment see Data S3). We then used maximum likelihood (Fig 2) and neighbor-joining (Fig S3) analyses to construct a phylogenetic tree based on this alignment. We were able to draw the same conclusions from both trees except for one major difference, which we discuss below. Both phylogenetic analyses were in agreement with expected branching topology of the *Drosophila* species [40] and concurred with our analyses of shared synteny (Fig 1A). For instance, *D. eugracilis Cid2* (clade A, orange branch) grouped with *Cid1* genes of the *melanogaster* group with high confidence. Its closest phylogenetic neighbor was the *Cid1* pseudogene from *D. eugracilis,* supporting *Cid2’s* species-specific origin in a recent ancestor of *D. eugracilis.* We also found that the *Cid1* and *Cid5* genes of the *Drosophila* subgenus form monophyletic sister clades (clade D is sister to clade E, Fig 2 and S3). We found that *D. busckii* and *D. albomicans* encode *Cid1* genes (clade E), based on phylogeny and shared synteny. However, whereas *D. albomicans* also encodes *Cid5, D. busckii* does not (clade D). The phylogenetic resolution between *Cid1* and *Cid5* clades is strong enough to suggest that the *Cid5* duplication may have predated the split between *D. busckii* and other members of the *Drosophila* subgenus, but that *Cid5* was subsequently lost in *D. busckii.*

**Fig 2.**
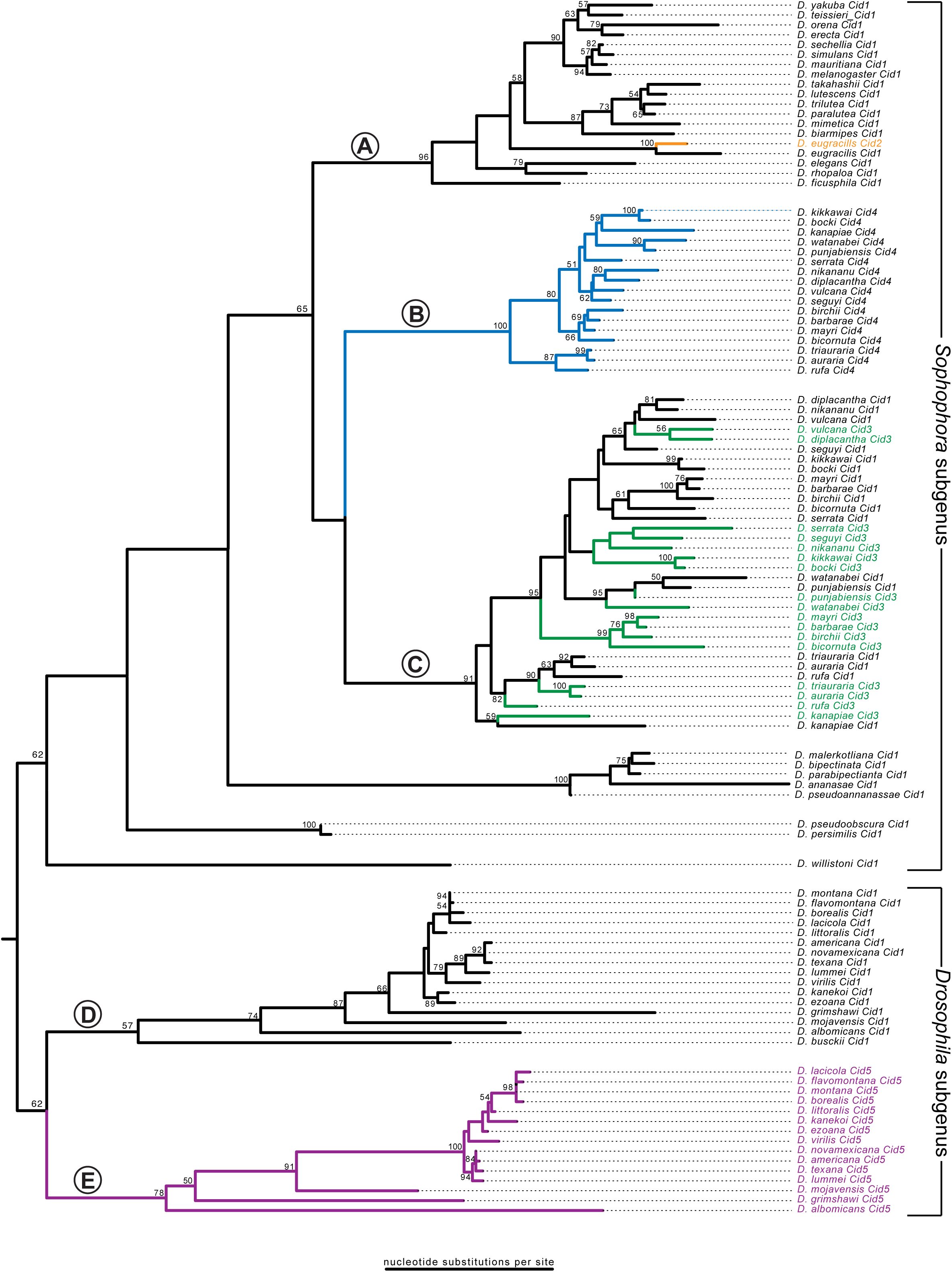
Evolutionary relationship among all *Drosophila Cid* paralogs. We performed maximum likelihood phylogenetic analyses using PhyML with a nucleotide alignment of the histone fold domain of all *Cid* paralogs. We found that *Drosophila* subgenus *Cid1* (clade E), *Drosophila* subgenus *Cid5* (clade D) and *montium* subgroup *Cid4* (clade B) all formed well-supported monophyletic clades suggesting a single origin for these *Cid* paralogs. In contrast, *montium* subgroup *Cid1* and *Cid3* grouped together (clade C), consistent with our finding that they may be undergoing recurrent recombination (Fig 3). Selected clades (labeled with letters A – E) are further discussed in the main text. Bootstrap values greater than 50 are shown. The tree is arbitrarily rooted to separate the *Sophophora* and *Drosophila* subgenera. Scale bar represents number of substitutions per site.

We also found that the *Cid4* genes from the *montium* subgroup form a monophyletic clade (Fig 2, clade B) that forms sister clade to the *montium* subgroup *Cid1* and *Cid3* genes (clade C). The *melanogaster* subgroup *Cid1* genes (clade A) formed an outgroup to *montium* subgroup genes *Cid1, Cid3* and *Cid4* (clade A is an outgroup to clade B and C). This was the only major difference in branching topology between the maximum likelihood and neighbor-joining analyses; the latter (Fig. S3) placed the *Cid4* genes from the *montium* subgroup (clade B) as a sister lineage to the *melanogaster* subgroup *Cid1* clade (clade A). Since Cid1 is expected to be the ancestral gene in both subgroups, we favor the tree topology suggested by the maximum likelihood analysis. Both analyses reveal an unexpected intermingling of the *montium* subgroup *Cid1/Cid3* genes into a single clade (Fig 2 & S3, clade C). This intermingled phylogenetic pattern could be the result of multiple, independent duplications of *Cid3* from *Cid1* in the *montium* subgroup. Alternatively, this pattern could reflect the effects of recurrent gene conversion, in which at least the HFD regions of *Cid1* and *Cid3* were homogenized by recombination.

Gene conversion between *Cid1* and *Cid3* could be facilitated by the close proximity of their genomic locations (see Fig 1A, ‘*Cid1* locus’ column), since frequency of gene conversion is inversely proportional to the distance between recombining sequences [42]. We used GARD (Genetic Analysis for Recombination Detection) analyses [43] to formally test for recombination between *Cid1* and *Cid3* from the *montium* subgroup. Consistent with our hypothesis of gene conversion, we found strong evidence for recombination between *Cid1* and *Cid3* (p = 0.0002) but not between *Cid1* and *Cid4.* The predicted recombination breakpoint is at the transition between the N-terminal tail and HFD domains (Fig 3A). Indeed, when we made a maximum likelihood tree from segment 1 alone (consisting primarily of the N-terminal tail), *Cid1* and *Cid3* formed the expected monophyletic clades distinct from each other (Fig 3B). However, when we made a maximum likelihood tree of the HFD, we found evidence for at least three specific instances of gene conversion (Fig 3C, recombination highlighted by asterisks). The HFD is important for Cid’s interaction with other nucleosome proteins as well as for centromere targeting [44-46]. We speculate that such a recombination pattern allows Cid1 and Cid3 to perform distinct functions due to their divergent N-terminal tails whereas the homogenization of the HFD ensures that both proteins retain function and localization to the centromeric nucleosome. This pattern of ancient divergence followed by recurrent gene conversion may also partially explain the discrepant phylogenetic position of the *Cid1/Cid3* clade from the *montium* subgroup relative to the *Cid4* clade from the same subgroup (compare Fig 2 to Fig S3).

**Fig 3.**
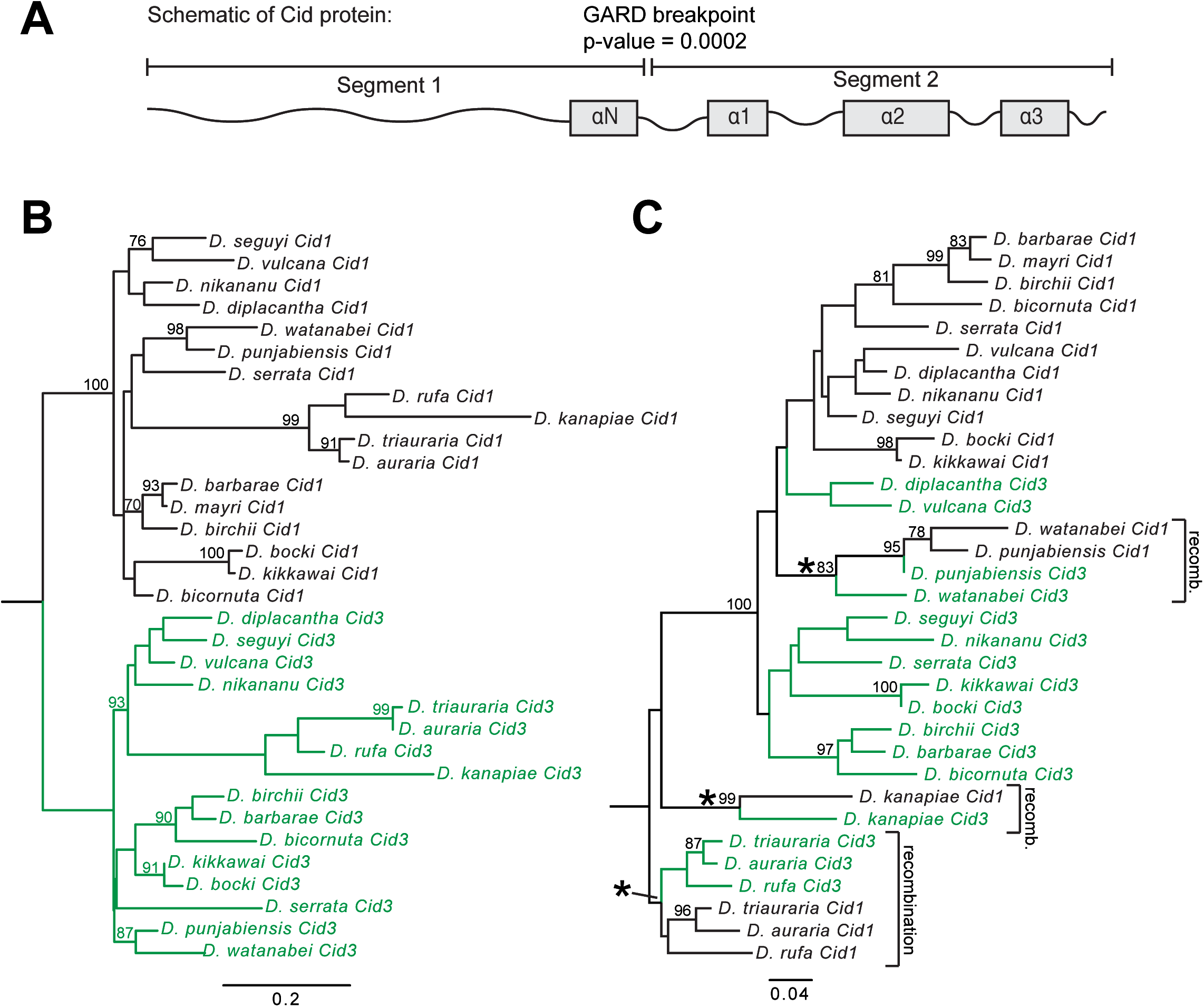
*Cid1* and *Cid3* have undergone recurrent gene conversion in the *montium* subgroup. (A) We used the Genetic Algorithm for Recombination Detection (GARD [43]) to test for recombination in the *montium* subgroup *Cid1* and *Cid3.* GARD identified one significant (p=0.0002) breakpoint between the N-terminal tail and the histone fold domain. (B, C) Maximum likelihood phylogenetic trees from an alignment of GARD segment 1 (B) and GARD segment 2 (C) were subsequently generated using PhyML. Bootstrap values above 75 are displayed. Asterisks indicate branches along which gene conversion likely occurred. Scale bar represents nucleotide substitutions per site.

### Drosophila Cid paralogs localize to centromeres

There are three possible outcomes following a functional gene duplication event: subfunctionalization, neofunctionalization and redundant functions, which often leads to the loss of one paralog. Because we observe the co-retention of most *Cid* duplicates for millions of years (with the exception of *Cid1* loss in *D. eugracilis* and *Cid3* loss in *D. mayri)*, it is unlikely that duplicate *Cid* genes have been retained for redundant functions. We therefore wanted to distinguish between the possibilities of subfunctionalization and neofunctionalization for duplicate *Cid* genes.

It is not unprecedented that a histone variant paralog might develop a new function. For example, in mammals, the H2B variant SubH2Bv acquired a non-nuclear role in acrosome development in sperm [47]. To assess the possibility that the *Cid* paralogs may have acquired a non-centromeric role *(i.e.,* have become neofunctionalized), we turned to cell biological analyses to determine their localization. Previous studies showed that Cid1 orthologs (including those from *D. kikkawai* and *D. virilis)* can fail to localize to *D. melanogaster* centromeres, due to changes at the interface between Cid1 and its chaperone protein CAL1 [48]. We therefore decided to test the localization of selected Cid paralogs in tissue culture cells from the same species.

Among all *montium* subgroup species that contain *Cid1*, *Cid3* and *Cid4,* cell lines were available only from *D. auraria* (cell line ML83-68, DGRC). We cloned the *Cid1*, *Cid3* and *Cid4* genes from *D. auraria* and tagged each with an N-terminal Venus tag to aid in visualization. We then transfected these constructs individually into *D. auraria* cells. We found that each Venus-Cid paralog localized in a similar manner, in punctate foci in a DAPI-intense region of the cells (Fig 4A). This pattern is highly characteristic of centromere localization [49]. To confirm this, we co-stained the cells with an antibody against CENP-C, a constitutively centromeric protein. Since no *D. auraria-specific* CENP-C antibodies were available, we first confirmed that the *D. melanogaster* CENP-C antibody appropriately marked centromeres in *D. auraria.* Indeed, the *D. melanogaster* CENP-C antibody recognized foci at the primary constriction of *D. auraria* metaphase chromosomes (Fig S4). Moreover, we found that Venus-Cid1, Venus-Cid3 and Venus-Cid4 all co-localized with CENP-C in this cell line (Fig 4A). Based on this, we conclude that all the *D. auraria* Cid paralogs localize to centromeres.

**Fig 4.**
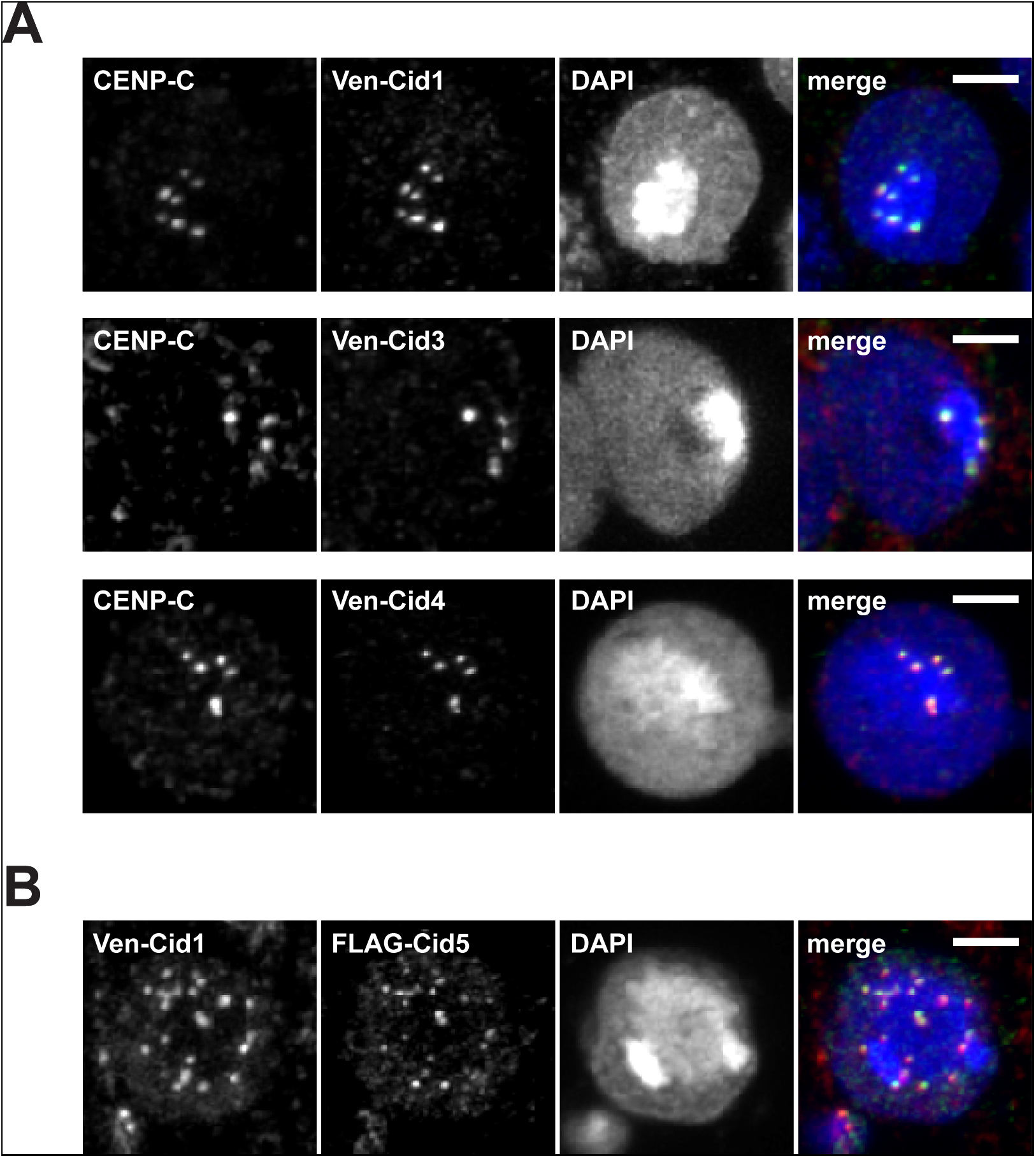
Proteins encoded by *Cid* paralogs localize to centromeres in cell culture. (A) Venus-tagged *D. auraria* Cid1, Cid3 *and* Cid4 were transiently transfected in a *D. auraria* cell line (top, middle and bottom panels, respectively). Cells were fixed and co-stained with a *D. melanogaster* CENP-C antibody (red in merged image) and anti-GFP (green in merged image). These data show co-localization of all three *montium* subgroup Cid proteins with CENP-C. (B) We co-transfected Venus-tagged Cid1 and FLAG-tagged Cid5 from *D. virilis* into a *D. virilis* cell line. Venus-Cid1 (red in merged image) and FLAG-Cid5 (green in merged image) both formed co-localized foci in the nucleus. All scale bars indicate a distance of two microns.

We similarly tested the localization of *D. virilis* Cid1 and Cid5 in a *D. virilis* cell line (WR Dv-1). Unfortunately, the antibody raised against *D. melanogaster* CENP-C did not recognize *D. virilis* centromeres likely due to the high divergence between the CENP-C orthologs from the two species. We therefore co-transfected Venus-Cid1 and FLAG-Cid5. We found that Cid1 and Cid5 co-localize at nuclear foci, in a staining pattern that is typical of centromeric localization (Fig 4B). Together, these results do not support the hypothesis that the simultaneous retention of multiple Drosophila *Cid* paralogs is due to non-centromeric function. Moreover, they suggest that despite their divergence, all Cid duplicates retain the ability to be recognized and deposited at centromeres by the existing CAL1-dependent machinery. Alternatively, Cid paralog proteins might achieve centromeric co-localization by forming heterodimers with Cid1. Based on these findings, we next investigated the possibility that subfunctionalization led to the simultaneous retention of all *Cid* paralogs by looking at tissue-specific expression.

### Testis restricted expression of *Cid3* and *Cid5*

One means by which subfunctionalization can occur is by tissue-specific expression [50, 51]. Duplicate genes could retain different subsets of promoter and enhancer elements from their parent gene, requiring both genes’ expression to fully recapitulate parental gene expression [52]. We therefore wondered whether any of the *Cid* duplicates showed tissue-specific expression. We expected that at least one *Cid* paralog in each species must have maintained mitotic function and would therefore be widely expressed in somatic tissues. To test this, we first looked for expression of *Cid* paralogs in *D. auraria* and *D. virilis* tissue culture cell lines, which are derived from embryonic and larval tissues, respectively. We extracted RNA from both cell lines and performed RT-PCR. After 30 cycles of PCR, we detected a faint *Cid1* band in addition to a robust *Cid4* band in the *D. auraria* cell line (Fig 5A). In the *D. virilis* cell line, we detected expression of *Cid1* but not *Cid5* after 30 cycles of PCRs (Fig 5B). We did not detect *Cid3 (D. auraria)* or *Cid5 (D. virilis)* in this assay, which suggests that both genes are either not expressed or are expressed at low levels in tissue culture cells. From this analysis, we predict that Cid4 (and possibly Cid1) performs somatic Cid function in *D. auraria (i.e.,* mitotic cell divisions for growth) and that Cid1 performs somatic Cid function in *D. virilis*.

**Fig 5.**
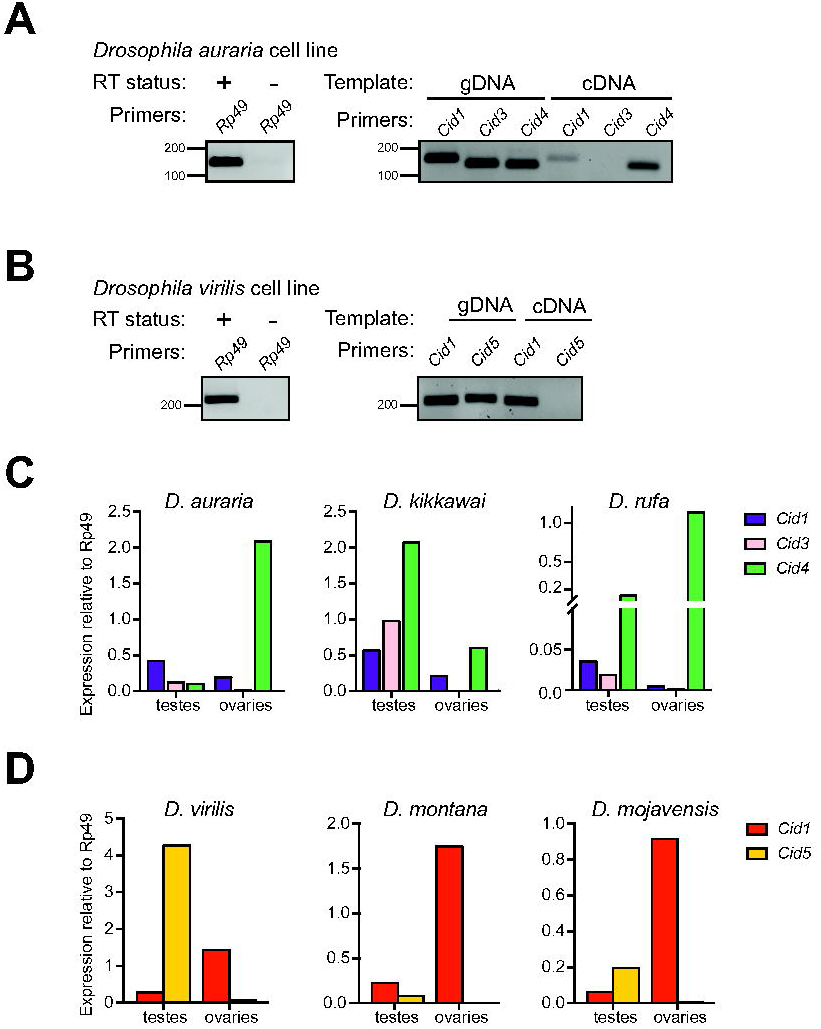
Male germline-restricted expression of some *Cid* paralogs. (A) Left gel: RNA samples used for *D. auraria* RT-PCR were free of DNA contamination as indicated by performing 35-cycle PCR for *Rp49* on cDNA samples generated with (+) and without (-) reverse transcriptase. Right gel: 30-cycle PCR performed with either genomic DNA (gDNA) or cDNA for *Cid1*, *Cid3* and *Cid4* from a *D. auraria* cell line. We detected both *Cid1* and *Cid4* expression but the *Cid4* expression band was more robust than the *Cid1* band. We did not detect expression of *Cid3* in this cell line. (B) Left gel: as in (A), RNA samples used for *D. virilis* RT-PCR were free of DNA contamination. Right gel: RT-PCR analyses of *Cid1* and *Cid5* from a *D. virilis* cell line at 30 cycles revealed only the expression of *Cid1.* We did not detect *Cid5* by RT-PCR. (C) RT-qPCR for *Cid1*, *Cid3* and *Cid4* from dissected tissues from three *montium* subgroup species revealed that *Cid1* and *Cid4* are expressed in both the testes and the ovaries whereas *Cid3* expression is testis restricted. (D) RT-qPCR from dissected tissues from three species from the *Drosophila* subgenus revealed that *Cid1* is expressed in the testes and ovaries of all three species whereas *Cid5* is only expressed in the testes. All RT-qPCR was normalized using *Rp49* as a control.

To further explore tissue specific expression, we performed RT-qPCR on dissected male and female *D. virilis* and *D. auraria* flies (whole fly, head, testes/ovaries, carcass). We performed the same analysis for *D. melanogaster,* which only encodes a single *Cid1* gene, for comparison. In *D. melanogaster,* we found that *Cid1* expression is highest in testes and ovaries and is relatively low in head and carcass (Fig S5A). This is not surprising since testes and ovaries contain higher numbers of actively dividing cells than the head and the carcass. Similarly, in *D. auraria* and *D. virilis*, we found low expression of *Cid* paralogs in the head and the carcass of male and female flies (Fig S5B, S5C). Interestingly, we found that the expression of *Cid3* in *D. auraria* and *Cid5* in *D. virilis* was restricted to the male germline (Fig 5C, Fig 5D). We also found that *Cid1* and *Cid4* in *D. auraria* as well as *Cid1* in *D. virilis* are expressed in both testes and ovaries.

We wanted to extend our expression analyses of the *Cid* paralogs to other species containing duplicate *Cid* genes. We performed RT-qPCR on two additional *montium* subgroup species (*D. kikkawai* and *D. rufa)* and on two additional *Drosophila* subgenus species (*D. montana* and *D. mojavensis).* In all cases, *Cid3* or *Cid5* expression was detected in testes but not in ovaries. *Cid1* and *Cid4* expression patterns were similar across species too, with the exception of *Cid1* in *D. rufa,* which expressed at very low levels in ovaries (Fig 5C, 5D, S4B, S4C).

Our findings are consistent with the hypothesis of tissue-specific specialization of the *Cid* paralogs in both the *montium* subgroup and the *virilis* group. These results also suggest that *Cid3* and *Cid5* were retained to perform a testis-specific function. In contrast, the other *Cid* paralogs are expressed in both somatic and germline tissues. However, these analyses lack the cellular resolution necessary to conclude whether the expression patterns are mutually exclusive or overlapping in tissues where multiple *Cids* are expressed. Moreover, in the *montium* subgroup, *Cid4* is expressed broadly in a pattern similar to *D. melanogaster Cid1,* and it is the primary *Cid* duplicate expressed in somatic cells. This suggests that *Cid4,* and not *Cid1,* performs canonical *Cid* function in *montium* subgroup species.

### Differential retention of N-terminal tail motifs and the evolution of new motifs following *Cid* duplication

Given their sequence divergence and different expression patterns, it seems likely that *Cid* paralogs may have subfunctionalized to perform distinct functions. Unlike the structural constraints that shape the HFD, the N-terminal tail of Cid is highly variable in length and sequence. We speculated that analyses of selective constraint in the N-terminal tail might present an additional opportunity to determine if subfunctionalization had occurred among the Cid paralogs. Although the specific function of the N-terminal tail has yet to be elucidated for *Drosophila* Cid, studies in humans and fission yeast have shown that the N-terminal tail is important for recruitment and stabilization of inner kinetochore proteins [22, 53, 54]. Furthermore, post-translational modifications of the N-terminal tail have been shown to be important for CENP-A mitotic function [55] and for facilitating interaction between two CENP-A molecules [56].

Conserved motifs provide an avenue to evaluate differential selective constraint in the N-terminal tail of different *Drosophila Cid* paralogs [57]. Motifs are regions of high similarity among protein sequences. They represent putative sites of protein-protein interaction and post-translational modification. We reasoned that we might be able to use the presence of certain N-terminal tail motifs as a proxy for various functional domains. We therefore used the motif generator algorithm, MEME [58], to identify conserved motifs in the N-terminal tail from six different groups of *Drosophila* Cid proteins: *melanogaster* group Cid1 (single copy genes only), *montium* subgroup Cid1, *montium* subgroup Cid3, *montium* subgroup Cid4, *virilis* group Cid1, and *virilis* group Cid5 (Fig S6). We then used the motif search algorithm, MAST [59], to search for each motif in all Cid proteins. In total we found 10 unique motifs (Fig S6). Finally, we overlaid our motif analysis with the Drosophila species tree to gain insight into the evolution of N-terminal tail motifs (Fig 6A).

**Fig 6.**
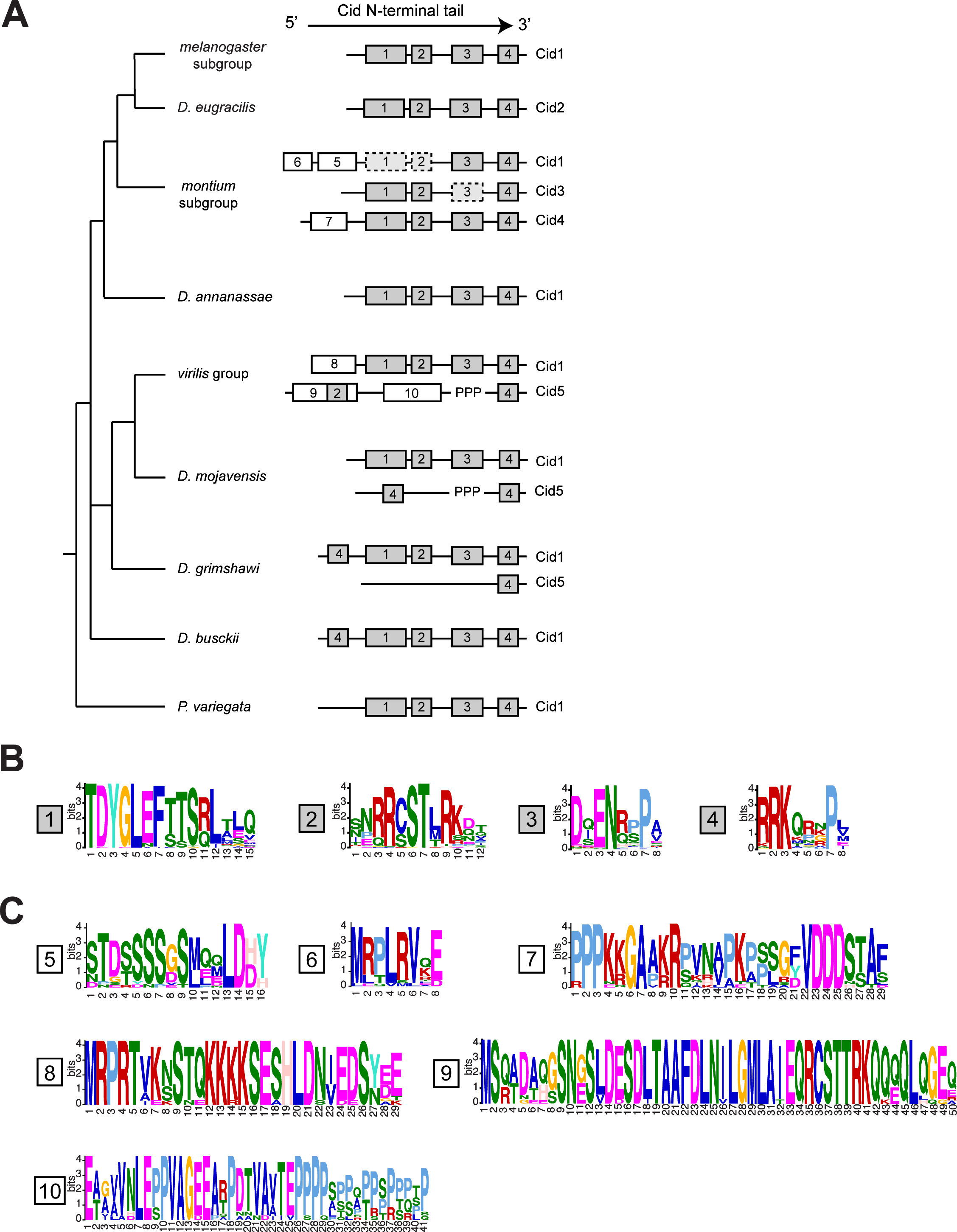
Evolution of N-terminal motifs among all Cid proteins. (A) A *Drosophila* species tree with a schematic of N-terminal tail motifs identified by MEME and MAST displayed to right of each species or species group. Each number represents a unique motif that does not statistically match any other motif in the figure with the exception of motif 2 and 9 (see methods). Gray boxes indicate ‘core’ motifs 1 – 4, which are present in all single copy *Cid* genes. White boxes indicate lineage specific motifs. ‘PPP’ indicates the position of the variable proline-rich region in Cid5. Dashed boxes indicate cases in which a given motif was present in ~50% of species. (B) Logos generated by MEME for consensus motifs 1 – 4. (C) Logos generated by MEME for consensus motifs 5 – 10.

From this analysis we can make several interesting conclusions. First, motifs 1-4 (Fig 6B) are conserved in every Cid1 protein when it is the only copy encoded in the genome. These motifs correspond nicely to the motifs we previously identified in the *melanogaster* group using Block Maker [41]. Although their function remains largely uncharacterized, motif 4 has been shown to be involved in recruitment of mitotic checkpoint protein, BubR1 [60]. Motif 4 could also play a role in histone-DNA interaction because it is located in the region where the N-terminal tail exits the nucleosome and passes between the two strands of DNA [45]. Motif 4 is the only motif present in all Cid paralogs, which suggests that it performs a general function among all Cids. Given their retention in all single copy Cid-containing Drosophila species, we consider motifs 1 – 4 to be the “core” Cid1 motifs (Fig 6B) and speculate that all are required for Cid1 function when it is the only centromeric histone protein. Indeed, all Drosophila species contain all of these motifs amongst their various Cid paralogs.

Next, we observed that some Cid paralogs had evolved and retained ‘new’ N-terminal tail motifs (Fig 6C). We identified three motifs that evolved in Cid paralogs from the *montium* subgroup; motifs 5 and 6 are found in Cid1 whereas motif 7 is found in Cid4. One might interpret the invention of additional N-terminal tail motifs as evidence of neofunctionalization, *e.g*. by invention and retention of novel protein-protein interactions. We also found that ‘ancestral’ motifs 1-3 appear to have been frequently lost from Cid1 and Cid3 whereas they are completely preserved in Cid4 (Fig 6A, dotted lines indicate motif is absent from ~50% of queried species). Intriguingly, some Cid1 and Cid3 orthologs in the *montium* subgroup appear to have differentially retained motifs 1-3; Cid1 has motif 3 and Cid3 has motifs 1 and 2. This differential retention of an ancestrally conserved subset of core motifs is suggestive of subfunctionalization [57]. Furthermore, our findings support the hypothesis that in the *montium* subgroup, it is the Cid4 paralog rather than the ancestral Cid1, which performs the canonical functions of centromeric histones carried out by Cid1 in other species, because Cid4 contains all core motifs but *montium* subgroup Cid1 does not. This would also be consistent with our expression analyses, in which Cid4 expresses more robustly than Cid1 in somatic cells (Fig 5A).

This pattern of new motif evolution and ancient motif degeneration is also evident in the Cid paralogs from the *virilis* group. In this group of species, the Cid1 paralog has retained the core set of motifs 1-4 but added motif 8. In contrast, Cid5 paralogs have added motifs 9 and10 but lost core motifs 1 and 3. We therefore conclude that the tissue-specific pattern of expression and the differential retention of N-terminal motifs support a general model of subfunctionalization, but that some paralogs may have acquired novel protein-protein interaction motifs perhaps to optimize for new, specialized centromeric functions.

### Different evolutionary forces act on different *Cid* duplicates

Tissue specific expression of some *Cid* paralogs and differential retention of N-terminal tail motifs supports the hypothesis that Cid paralogs may have subfunctionalized. We therefore considered the possibility that duplicate *Cid* genes were retained to allow optimization for divergent functions. In the *melanogaster* group, *Cid1* (a single copy *Cid* gene) has been shown to evolve rapidly [7], perhaps due to its interaction with rapidly evolving centromeric DNA and the need for drive suppressors in male meiosis [17]. While this rapid evolution might be required for the ‘drive suppressor’ function, it may be disadvantageous for canonical Cid function *(e.g.,* mitosis). As a result, selection may act differently on *Cid* in the male germline than on somatic or ovary-expressed *Cid.* For instance, some *Cid* paralogs *(e.g.,* those that are expressed primarily in the male germline and may suppress centromere-drive) might evolve under positive selection while others would not.

We used maximum likelihood methods using the PAML suite to test for positive selection on each of the *Cid* paralogs. For *montium* subgroup *Cid1* and *Cid3,* we performed each analysis separately on GARD segment 1 and 2 (Fig 3). For all other *Cid* genes we performed PAML analyses on full-length alignments (Data S4, Data S5). Consistent with our prediction, we found that some, but not all, *Cid* paralogs evolve under positive selection (Fig 7A). For example, PAML analyses reveal that *Cid3* segment 1 evolved under positive selection (Table S2, p = 0.01). However, we did not find evidence that *Cid5,* another male germline-restricted paralog, evolves under positive selection. We note, however, that we were unable to unambiguously align a highly variable proline-rich segment in Cid5’s N-terminal tail and excluded this segment from our analyses (Fig 7B). If positive selection was occurring in this region, we would be unable to detect it. We also found that *Cid4* evolved under positive selection but *montium* subgroup *Cid1* and *Cid3* segment 2, and *virilis* group *Cid1,* did not (Fig 7A, Table S2).

**Fig 7.**
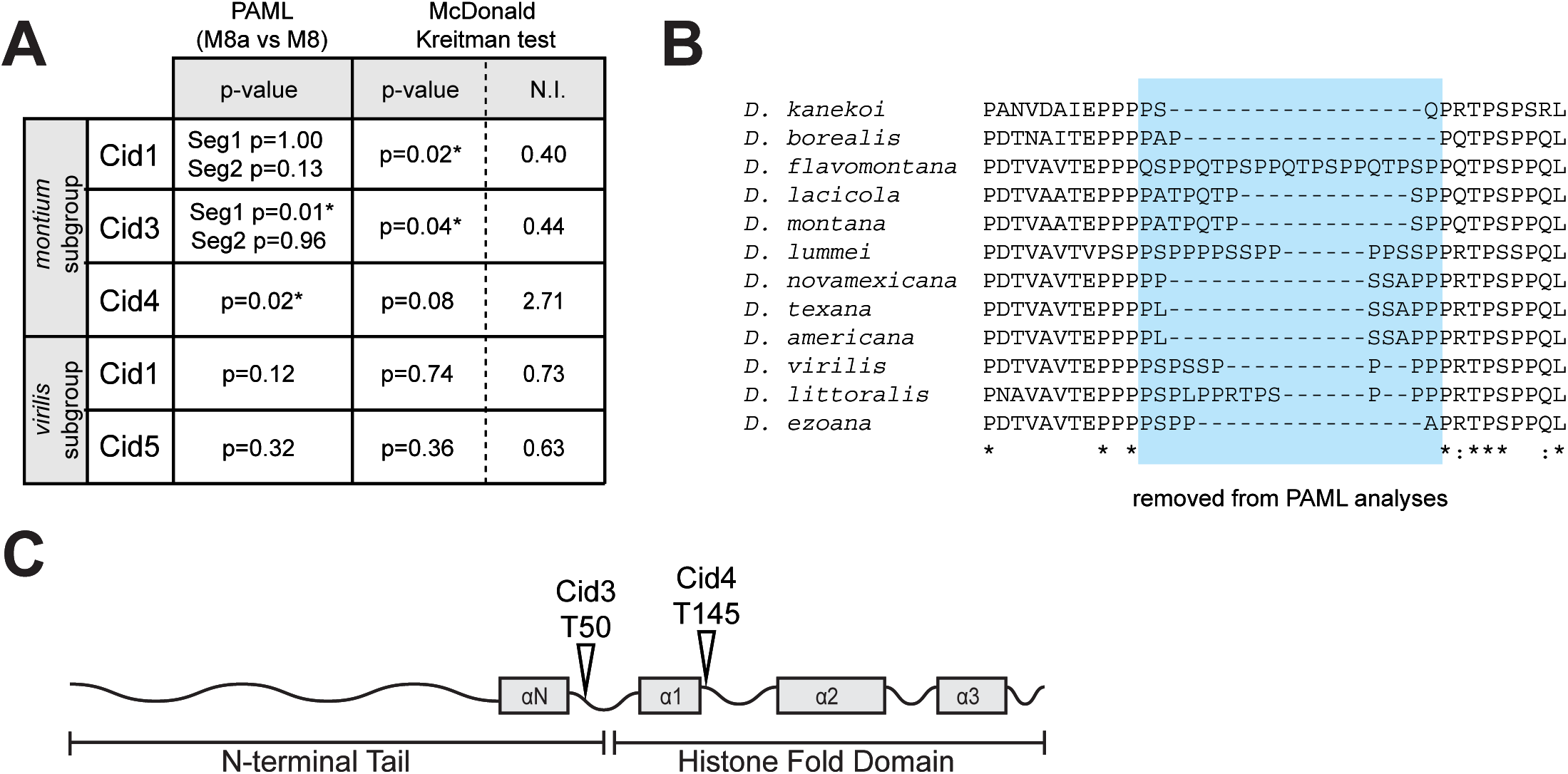
Different Cid paralogs evolve under different evolutionary pressures. (A) Summary of tests for positive selection performed on each *Cid* paralog. Tests that were statistically significant (p<0.05) are indicated with an asterisk. For the McDonald Kreitman test, Neutrality Index (N.I.) is also displayed. N.I. < 1 indicates an excess of non-synonymous fixed differences between species and suggests positive selection. N.I. > 1 indicates fewer nonsynonymous fixed differences than expected and suggests purifying selection. (B) A protein alignment of Cid5 from *virilis* group species. The variable, proline-rich region which was excluded from PAML tests for positive selection is highlighted in blue. (C) A schematic of a representative Cid protein, showing sites evolving under positive selection identified by Bayes Emperical Bayes analyses (posterior probability > 0.95).

For those genes that PAML identified as having evolved under positive selection *(Cid3* segment 1 and *Cid4),* Bayes Empirical Bayes analyses identified one amino acid in Cid3 and one amino acid in Cid4 as having evolved under positive selection with a high posterior probability (>0.95). In Cid3, the positively selected site is adjacent to the αN-helix. In Cid4, the positively selected site is in loop 1 of the HFD (Fig 7C, Table S2). Interestingly, these are both places where Cid is predicted to contact centromeric DNA [45]. These results are consistent with the hypothesis that both *Cid3* and *Cid4* are engaged in a genetic conflict involving centromeric DNA.

We next used the McDonald-Kreitman (MK) test to look for positive selection in each of the *Cid* paralogs. While PAML detects positive selection occurring recurrently at selected amino acid residues across deep evolutionary time, the MK test detects more recent positive selection distributed over entire genes or protein domains. For the *montium* subgroup, we sequenced and compared *Cid1*, *Cid3* and *Cid4* paralogs from 26 strains of *D. auraria* and 10 strains of *D. rufa.* For *virilis* group *Cids,* we sequenced *Cid1* and *Cid5* paralogs from 10 strains of *D. virilis* and 21 strains of *D. montana* (Data S6, Data S7). We found an excess of non-synonymous fixed differences between *D. auraria* and *D. rufa Cid1* and *Cid3,* suggesting that both genes evolve under positive selection (Fig 7A, Table S3). Parsing the signal by performing the MK test on just the N-terminal tail or just the HFD domain revealed that *Cid1* and *Cid3* HFD domains evolve under positive selection (Table S3). However, we did not find evidence for positive selection in the N-terminal tails. Most of the non-synonymous fixed differences occur in Loop1, which is predicted to contact centromeric DNA [45]. Interestingly, even though PAML analyses detected ancient recurrent positive selection in *montium* group *Cid4,* we found no evidence for more recent positive selection since the *D. auraria-D. rufa* divergence using the MK test (p=0.08, Neutrality Index = 2.71). We also found no evidence of positive selection having acted on *virilis* group *Cid1* or *Cid5* using the MK test (Fig 7A, Table S3).

To summarize our positive selection analyses, we found that *Cid3* has experienced both ancient and recent positive selection in protein domains predicted to contact centromeric DNA. *Cid4* also has also experienced ancient, recurrent positive selection at putative DNA-contacting sites, but we found no evidence of recent positive selection in a MK test comparison. This could suggest that *Cid4* was either relieved of its role in such conflict or that the MK test lacks the power to detect selection acting on only a few residues. Similarly, although PAML analyses failed to identify a pattern of ancient, recurrent positive selection, the MK test did reveal positive selection for *montium* subgroup *Cid1* while comparing the entire HFD. In contrast, we did not find evidence for positive selection having acted on *Cid1* and *Cid5* in the *virilis* group by either test.

### Discussion

The prevailing model that centromeric histones are encoded by single copy genes likely stems from the fact that model organisms like *D. melanogaster* have only one copy [12]. Our investigation into *Cid* duplications in *Drosophila* has more than doubled the number of known *CenH3* duplications in animals. Based on our detailed analysis of two subgenera *(Drosophila* and *Sophophora),* we predict that over one thousand *Drosophila* species encode two or more *CenH3 (Cid)* genes [61]. We further conclude that *D. melanogaster* and other *Drosophila* species that have only one *Cid* are the minority; most *Drosophila* species have multiple *Cid* paralogs.

The availability of many high-quality sequenced genomes as well as the comprehensive understanding of phylogenetic relatedness between species make *Drosophila* an ideal system to study gene duplication and evolution. This facilitated our discovery of four ancient *Cid* duplications in *Drosophila.* We found that while *Cid1* (previously known as just *‘Cid’)* is consrved in its shared syntenic location in all species examined except one, many species encode one or two additional *Cid* genes. The species of the *montium* subgroup, including *D. kikkawai,* have three *Cid* genes *(Cid1, Cid3 and Cid4),* which were born from a duplication event ~15 million years ago. The species of the *virilis* group, as well as *D. mojavensis* and *D. grimshawi (repleta* and *Hawaiian* groups, respectively), have two *Cid* genes *(Cid1* and *Cid5)*, which were born from a duplication event ~40 million years ago. These *Cid* duplications have been almost completely preserved in extant species. Despite the fact *Cid* paralogs are divergent from one another at the sequence level, all paralogs have the ability to localize to centromeres when expressed in tissue culture cells.

Our phylogenetic analyses support our synteny-based conclusions, and reveal recurrent recombination between *Cid1* and *Cid3* in *montium* subgroup species. This is the first reported case of recombination between *CenH3* paralogs. Our results suggest that this recombination results in evolutionary homogenization of the histone fold domain between *Cid1* and *Cid3,* while the N-terminal tails of *Cid1* and *Cid3* appear to be evolving independently, perhaps maintaining divergent functions. This recombination could be the genetic mechanism by which *Cid1* and *Cid3* maintain function in the centromeric nucleosome via near-identical HFDs despite having divergent N-terminal tails, which facilitates distinct interactions. This pattern of gene conversion is akin to patterns of recombination seen for paralogous mammalian antiviral proteins, IFIT1 and IFIT1B, in which gene conversion homogenizes the N-terminal oligomerization domain but not the divergent C-terminus, which allows IFIT1 and IFIT1B proteins to have distinct anti-viral specificities [62].

What is the evidence that *Cid* paralogs have distinct functions? The strongest evidence is that they have been co-retained in both the *montium* subgroup and the virilis/repleta/Hawaiian radiation for tens of millions of years. If they performed redundant functions, we predict that one of the paralogs would be lost over this time frame considering the high rate of DNA deletion in *Drosophila* [63]. Indeed, we observed only two instances of Cid duplication followed by pseudogenization *(Cid3* pseudogene in *D. mayri* and *Cid1* pseudogene in *D. eugracilis)* and inferred the possible loss of *Cid5* (in *D. busckii)*. Our finding that *Cid3* and *Cid5* are expressed primarily in the male germline also supports our subfunctionalization hypothesis. Finally, differential retention of N-terminal tail motifs and different selective pressures on different *Cid* genes further supports the idea that these *Cid* paralogs do not have redundant roles.

Interestingly, our expression and motif analyses strongly suggest that *Cid4* has taken over the primary function of somatic centromeric histone function in *montium* subgroup species. *Cid4* is the primary *Cid* gene expressed in *D. auraria* tissue culture cells and is the only Cid paralog in this species that contains all four of the ‘core’ N-terminal tail motifs. In contrast, the ‘ancestral’ *Cid1* is expressed at lower levels than *Cid4, Cid3* is primarily expressed in the male germline, and neither *Cid1* nor *Cid3* contain all four ‘core’ motifs. This finding has implications for future experiments taking an evolutionary approach to study Cid function. The correct *Cid* paralog for such studies must be chosen carefully. Further functional experimentation, such as creating genetic knockouts, will be required to determine the specific function of each *Cid* paralog.

We propose that in species with a single-copy Cid gene, the same protein must perform multiple functions including mitotic cell division in somatic tissues and drive suppression in the male germline. These functions might require different selective pressures to achieve functional optimality. For example, we have previously proposed that drive suppression results in rapid evolution of *Cid* to co-evolve with rapidly evolving centromeric DNA [17] whereas mitotic function might impose purifying selection on *Cid,* minimizing changes in amino acid sequence. Therefore, it could be advantageous to have two copies of *Cid* such that each encodes a separate function. Our results suggest that *Cid3* and *Cid5* are candidate drive suppressors given their male germline-restricted expression. Consistent with this prediction, we detected evidence for positive selection in *Cid3.* In contrast, we did not find evidence that *Cid5* evolves under positive selection. This leaves open the possibility that *Cid5* performs an alternative, centromeric, male germline function independent of potential centromere-drive suppression in meiosis.

If it is advantageous to have multiple *Cid* paralogs, why don’t more animal species possess more than one gene encoding centromeric histones? We hypothesize that retention of duplicate *Cid* genes requires a defined series of evolutionary events and that the cadence of the mutations determines the ultimate fate of the duplicated genes [64]. First, the duplication must not be instantaneously harmful; gene expression must be carefully controlled, as Cid overexpression or expression at the wrong time during the cell cycle can be catastrophic [65, 66]. Even though other kinetochore proteins might limit Cid incorporation into ectopic sites [67], a duplicate *Cid* gene that acquired a strong or constitutive promoter would almost certainly be detrimental. Furthermore, in order for a duplicate *Cid* gene to be retained, a series of subfunctionalizing mutations must occur (before pseudogenization of either paralog) such that both paralogs are required for complete Cid function. This model, known as duplication-degeneration-complementation [51], most often refers to mutations in the promoters of duplicate genes. However, the same principle could be applied to mutations in coding regions. Since it is easier to introduce a mutation that results in a non-functional *Cid* gene than a subfunctionalized *Cid,* most *Cid* duplicates probably succumb to pseudogenization early in their evolutionary history and, in *Drosophila,* are quickly lost from the genome [68].

The existence of *Cid* duplications in genetically tractable organisms provides an opportunity to study the multiple functions of a gene that is essential when present in a single copy. While we know a lot about the role of *Cid* in mitosis, its roles in meiosis [69] and inheritance of centromere identity through the germline [70] are less well-characterized. Studying subfunctionalized *Cid* paralogs may allow for detailed analysis of these underappreciated *Cid* functions without the risk of disrupting essential mitotic functions. Future functional studies can now leverage the insight provided by duplicate *Cid* genes, where evolution and natural selection may have already carried out a ‘separation of function’ experiment.

## Materials and Methods

### *Drosophila* species and strains

Flies were obtained from the *Drosophila* Species Stock Center at UC-San Diego (https://stockcenter.ucsd.edu) and from the *Drosophila* Stocks of Ehime University in Kyoto, Japan (https://kyotofly.kit.jp/cgi-bin/ehime/index.cgi). For a complete list of species and strains used in this study, see Table S4.

### Identification of *Cid* orthologs and paralogs in sequenced genomes

*Drosophila Cid* genes were identified in previously sequenced genomes using both *D. melanogaster Cid1* and *H3* histone fold domain to query the non-redundant database using tBLASTn [71] implemented in Flybase [72] or NCBI genome databases. Since *Cid* is encoded by a single exon in *Drosophila,* we took the entire open reading frame for each *Cid* gene hit. For annotated genomes, we recorded the syntenic locus (3-prime and 5-prime neighbor genes) of each *Cid* gene hit as indicated by the Flybase genome browser track. For genomes that were sequenced but not annotated (*D. eugracilis*, *D. takahashii*, *D. ficusphila*, *D. kikkawai* and *P. variegata)*, we used the 3-prime and 5-prime nucleotide sequences flanking the putative *Cid* open reading frame as a query to the *D. melanogaster* genome using BLASTn. We annotated the syntenic locus according to these *D. melanogaster* matches. Each *Cid* gene was named according to its shared syntenic location. It is worth noting that the Flybase gene prediction for *D. virilis Cid5* (GJ21033) includes a predicted intron but we found no evidence that *Cid5* was spliced in any tissue (data not shown). The results of all BLAST searches are summarized in Table S1.

### Identification of *Cid* orthologs and paralogs in non-sequenced genomes

Approximately 10 whole (5 male, 5 female) flies were ground in DNA extraction buffer (10mM Tris pH 7.5, 10mM EDTA, 100mM NaCl, 0.5% SDS) with Proteinase K (New England Biolabs). Ground flies were incubated for 2 hrs at 55°C. DNA was extracted using phenol-chloroform (Thermo Fisher Scientific) according to the manufacturers instructions. Primers were designed to amplify each *Cid* paralog based on regions of homology in neighboring genes or intergenic regions. Only *Cid* paralogs that were predicted to be present in the species based on related species sequenced genomes were amplified. All PCRs were performed using Phusion DNA Polymerase (New England Biolabs). Appropriately sized amplicons were gel isolated and cloned into the cloning/sequencing vector pCR-Blunt (Thermo Fisher Scientific) and Sanger sequenced with M13F and M13R primers plus additional primers as needed to obtain sufficient coverage of the locus. A complete list of primers used in this study can be found in Table S5. A list of primer pairs used to amplify *Cid* paralogs in non-sequenced genomes can be found in Table S6. A list of Genbank accession numbers can be found in Table S4

### Phylogenetic analyses

*Cid* sequences were aligned using the ClustalW [73] ‘translation align’ function in the Geneious software package (version 6) [74]. Alignments were further refined manually, including removal of gaps and poorly aligned regions. Maximum likelihood phylogenetic trees of *Cid* nucleotide sequences were generated using the HKY85 substitution model in PhyML, implemented in Geneious, using 1000 bootstrap replicates for statistical support. Neighbor-joining trees correcting for multiple substitutions were generated using CLUSTALX [73]. We used the GARD algorithm implemented at datamonkey.org to examine alignments for evidence of recombination [43]. Pairwise percent identity calculations were made in Geneious. Phylogenies were visualized using FigTree (http://tree.bio.ed.ac.uk/software/figtree/) or Dendroscope [75]

### Cloning Cid fusion proteins

*Cid* genes from *D. auraria (Cid1, Cid3* and *Cid4)* and *D. virilis (Cid1* and *Cid5)* were amplified from genomic DNA and cloned into pENTR/D-TOPO (ThermoFisher). We used LR clonase II (ThermoFisher) to directionally recombine each *Cid* gene into a destination vector from the Drosophila Gateway Vector Collection, generating either N-terminal Venus (pHVW) or 3XFLAG (pHFW) fusion under the control of the *D. melanogaster* heat-shock promoter.

### Cell culture

Cell lines (*D. auraria* cell line ML83-68 and *D. virilis* cell line WR DV-1) were obtained from the Drosophila Genomics Resource Center in Bloomington, Indiana (https://dgrc.bio.indiana.edu). *D. auraria* cells were grown at room temperature in M3+BPYE + 12.5%FCS and *D. virilis* cells were grown in M3+BPYE + 10%FCS.

### Transfection experiments

2 micrograms plasmid DNA was transfected using Xtremegene HP transfection reagent (Roche) according to the manufacturer’s instructions. Cells were heat-shocked at 37°C for one hour 24 hours after transfection to induce expression of the *Cid* fusion protein.

### Imaging

Cells were transferred to a glass coverslip 48 hours after heatshock. Cells were treated with 0.5% sodium citrate for 10 min and then centrifuged on a Cytospin III (Shandon) at 1900rpm for 1 min to remove cytoplasm. Cells were fixed in 4% PFA for 5 min and blocked with PBSTx (0.3% Triton) plus 3% BSA for 30 minutes at room temperature. Coverslips with cells were incubated with primary antibodies at 4°C overnight at the following concentrations: mouse anti-FLAG (Sigma F3165) 1:1000, chicken anti-GFP (Abcam AB13970) 1:1000, rabbit anti-CENP-C (gift from Aaron Straight) 1:1000. Coverslips with cells were incubated with secondary antibodies for 1 hour at room temperature at the following concentrations: goat anti-rabbit (Invitrogen Alexa Fluor 568, A-11011) 1:2000, goat anti-chicken (Invitrogen Alexa Fluor 488, A-11039) 1:5000, goat anti-mouse (Invitrogen Alexa Fluor 568, A-11031) 1:2000. Images were acquired from the Leica TCS SP5 II confocal microscope with LASAF software.

### Expression analyses

RNA was extracted from *D. auraria* cell line ML83-68 and *D. virilis* cell line WR DV-1 using the TRIzol reagent (Invitrogen) according to the manufacturers instructions. To investigate expression profiles in adult tissues, RNA was extracted from whole bodies, and dissected tissues (heads, germline and the remaining carcasses) from *D. auraria, D. rufa, D. kikkawai, D. virilis, D. montana* and *D. mojavensis* flies. All samples were DNase treated (Ambion) and then used for cDNA synthesis (SuperScript III, Invitrogen). During cDNA synthesis, a ‘No RT’ control was generated for each RNA extraction in which the reverse transcriptase was excluded from the reaction. For RT-PCR experiments, the presence of genomic DNA contamination was ruled out by performing PCR that amplified the housekeeping gene, *Rp49,* on each cDNA sample as well as each ‘No RT’ control. 25- (not shown) and 30-cycle PCRs were performed with primers specific to each *Cid* paralog and samples were run on an agarose gel for visualization. RT-qPCR was performed according to the standard curve method using the Platinum SYBR Green reagent (Invitrogen) and primers designed to each *Cid* paralog and to *Rp49.* Reactions were run on an ABI QuantStudio 5 qPCR machine using the following conditions: 50°C for 2 min, 95°C for 2 min, 40 cycles of (95°C for 15s, 60°C for 30s). We ensured that all primer pairs had similar amplification efficiencies using a dilution series of genomic DNA. Three technical replicates were performed for each cDNA sample. Transcript levels of each gene were normalized to *Rp49.* For all primers used in RT-PCR and RT-qPCR experiments, see Table S5 and Table S6.

### Motif analyses

Motifs were identified in six different groups of Cid proteins (Fig S6) using the motif generator algorithm MEME [58] implemented on http://meme-suite.org/ [76]. Several motifs identified in different groups were similar to one another. For example, the motif “TDYLEFTTS” appeared in *melanogaster* group Cid1s, *montium* subgroup Cid3s and Cid4s and *virilis* group Cid1s (Fig S6, underlined residues). To determine which motifs were the same, we used the motif search algorithm MAST [59] to search for the top four motifs from each group against all 86 sequences used for motif generation. In total, we found 10 unique motifs (Fig 6B, 6C, S5). The only instance in which the motifs were not totally independent was for motif 2 and motif 9. Motif 2 was contained within motif 9, but motif 9 was significantly longer than motif 2 so we considered it to be an independent motif. We mapped all 10 motifs to the *Cid* genes in the six groups plus *D. eugracilis* Cid2, *D. mojavensis* and *D. grimshawi Cid1* and *Cid5, D. busckii,* and the outgroup species *P. variegata* Cid1. We considered a motif to be present in a given protein if the MAST p-value was < 10^−5^.

### Positive selection analyses

We used the PAML suite of programs [77] to test for positive selection on each *Cid* paralog across deep evolutionary time. Alignments for each *Cid* paralog were generated and manually refined as described above. Alignments (Data S5) and *Cid* gene trees were used as input into the CODEML NSsites model of PAML. To determine whether each *Cid* paralog evolves under positive selection, we compared two models that do not allow dN/dS to exceed 1 (M7 and M8a) to a model that allows dN/dS > 1 (M8). Positively selected sites were classified as those sites with a M8 Bayes Empirical Bayes posterior probability > 95%. We used the McDonald Kreitman (MK) test [78] to look for more recent positive selection at the population level. To implement the MK test for *montium* subgroup *Cid* paralogs we compared *Cid* sequences in 26 strains of *D. auraria* to 10 strains of *D. rufa.* In the *virilis* group, we compared *Cid* sequences in 10 strains of *D. virilis* to 20 strains of *D. montana.*

---

**S1 Fig. *Cid1* pseudogene in *D. eugracilis***

*D. eugracilis* genomic sequence at the *Cid1* syntenic locus. The coding sequence for *Cid1* neighbor genes, *crowded by cid* and *bb in a box car,* are highlighted. *Cid1* pseudogene sequence is underlined (dashed red line). An early stop codon is indicated with a red hexagon.

**S2 Fig. *Cid3* pseudogene in *D. mayri***

*D. mayri* genomic sequence at the *Cid1* locus as identified by PCR and Sanger sequencing. DNA sequence from an intron of the *Arrow* gene is highlighted in blue. *Cid3* pseudogene sequence is underlined (dashed blue line). An early stop codon is indicated with a red hexagon.

**S3 Fig. Neighbor-joining phylogenetic tree of *Drosophila Cid* genes**

We performed neighbor joining phylogenetic analyses with a nucleotide alignment of the histone fold domain of all *Cid* paralogs. We found that *Drosophila* subgenus *Cid1* (clade E), *Drosophila* subgenus *Cid5* (clade D) and *montium* subgroup *Cid4* (clade B) all formed well-supported monophyletic clades suggesting a single origin for these *Cid* paralogs. In contrast, *montium* subgroup *Cid1* and *Cid3* grouped together (clade C), consistent with our finding that they may be undergoing recurrent recombination (Fig 3). Selected clades (labeled with letters A – E) are further discussed in the main text. Bootstrap values greater than 50 are shown. The tree is arbitrarily rooted to separate the *Sophophora* and *Drosophila* subgenera. Scale bar represents number of substitutions per site.

**S4 Fig. *D. melanogaster* CENP-C staining in *D. auraria* tissue culture cells**

(A) *D. auraria* cell line ML83-68 was fixed and stained with *D. melanogaster* anti-CENP-C (red in merged image) and imaged using a confocal microscope. DNA (blue in merged image) shows metaphase chromosomes. (B) As in (A) except DNA shows interphase chromosomes.

**S5 Fig. Tissue specific expression of *Cid* paralogs, full panel**

(A) RT-qPCR from dissected tissues from *D. melanogaster Cid1.* (B) RT-qPCR for *Cid1, Cid3* and *Cid4* from dissected tissues from three *montium* subgroup species. *Cid1* and *Cid4* are expressed in both the testes and the ovaries whereas *Cid3* expression is testis-restricted. (C) RT-qPCR from dissected tissues from three *Drosophila* subgenus species. *Cid1* is expressed in the testes and ovaries of all three species but *Cid5* is only expressed in the testes. In all cases, *Cid* expression in the head and carcass is very low or undetectable.

**S6 Fig. MEME identified motifs**

List of motifs in logo format identified by MEME for Cid from six sequence groups. E-value and motif length is displayed to the right of each motif. An example of a motif that was found in several groups (“TDYLEFTTS”) is underlined.

**S1 Table. BLAST results**

List of all *Cid-*like hits from tBLASTn searches

**S2 Table. PAML results**

Summary table of M7 vs. M8 and M8a vs. M8 PAML results for each *Cid* gene or gene segment. P-values less than 0.05 are indicated in bold text. PP=posterior probability.

**S3 Table. McDonald-Kreitman test results**

Summary of results from McDonald-Kreitman tests for each *Cid* gene or gene segment. P-values less than 0.05 are indicated in bold text. Neutrality index > 1 is indicative of an excess of non-synonymous polymorphisms, which suggests negative selection. Neutrality index < 1 is indicative of an excess of non-synonymous fixed changes, which suggests positive selection.

**S4 Table. A list of *Drosophila* species and strains used in this study**

Table includes accession numbers for all *Cid* genes sequenced in this study.

**S5 Table. Primer Sequences**

List of primer sequences used in this study.

**S6 Table. Primer Pairs**

List of primer pairs used for various aspects of this study including: identifying *Cid* in species without sequenced genomes, sequencing *Cid* in strains of *D. auraria, D. rufa, D. virilis* and *D. montana,* RT-PCR, and RT-qPCR.

**S1 Data. *D. eugracilis Cid1* and *Cid2* sequences**

**S2 Data. Cid sequences for phylogenetic analyses**

Untrimmed sequences used to generate HFD alignments for maximum likelihood and neighbor-joining trees.

**S3 Data. Cid histone fold domain alignment**

Alignment used to generate maximum likelihood and neighbor-joining trees.

**S4 Data. Cid sequences for maximum likelihood selection analyses**

Untrimmed sequences used in PAML

**S5 Data. Cid alignments for PAML**

Trimmed sequence alignments used for PAML.

**S6 Data. Cid sequences used in MK Test**

Untrimmed sequences used in MK Tests.

**S7 Data. Alignments used in MK Test**

Trimmed sequence alignments used in MK Tests.

## Acknowledgements

We thank Rick McLaughlin, Antoine Molaro, Courtney Schroeder, Janet Young, Tera Levin and Rini Kasinathan for their comments on the manuscript and past and present members of the Malik lab for valuable discussions. We thank Frances Welsh and Tobey Casey for help with the PCR analyses to confirm the presence or absence of potential Cid paralogs. We thank the San Diego and Ehime Species stock centers for the use of Drosophila strains, and Aaron Straight for sharing the Drosophila CENP-C antibody.

